# Dynamics of corticospinal motor control during overground and treadmill walking in humans

**DOI:** 10.1101/177915

**Authors:** Luisa Roeder, Tjeerd W Boonstra, Simon S Smith, Graham K Kerr

## Abstract

Increasing evidence suggests cortical involvement in the control of human gait. However, the nature of corticospinal interactions remains poorly understood. We performed time-frequency analysis of electrophysiological activity acquired during treadmill and overground walking in 22 healthy, young adults. Participants walked at their preferred speed (4.2, SD 0.4 km h^−1^), which was matched across both gait conditions. Event-related power, corticomuscular coherence (CMC) and inter-trial coherence (ITC) were assessed for EEG from bilateral sensorimotor cortices and EMG from the bilateral tibialis anterior (TA) muscles. Cortical power, CMC and ITC at theta, alpha, beta and gamma frequencies (4-45 Hz) increased during the double support phase of the gait cycle for both overground and treadmill walking. High beta (21-30 Hz) CMC and ITC of EMG was significantly increased during overground compared to treadmill walking, as well as EEG power in theta band (4-7 Hz). The phase spectra revealed positive time lags at alpha, beta and gamma frequencies, indicating that the EEG response preceded the EMG response. The parallel increases in power, CMC and ITC during double support suggest evoked responses at spinal and cortical populations rather than a modulation of ongoing corticospinal oscillatory interactions. The evoked responses are not consistent with the idea of synchronization of ongoing corticospinal oscillations, but instead suggest coordinated cortical and spinal inputs during the double support phase. Frequency-band dependent differences in power, CMC and ITC between overground and treadmill walking suggest differing neural control for the two gait modalities, emphasizing the task-dependent nature of neural processes during human walking.

**New & Noteworthy:** We investigated cortical and spinal activity during overground and treadmill walking in healthy adults. Parallel increases in power, CMC and ITC during double support suggest evoked responses at spinal and cortical populations rather than a modulation of ongoing corticospinal oscillatory interactions. These findings identify neurophysiological mechanisms that are important for understanding cortical control of human gait in health and disease.

## Introduction

The control of human locomotion has commonly been thought to be driven by spinal and subcortical neural circuits. This assumption has been primarily based on animal studies, which show that cortical networks are only involved in generating locomotor activity in animals during more demanding walking tasks, such as precision stepping or obstacle avoidance (Armstrong 1988; Drew et al. 2008; Grillner 1985). Increasing evidence from human neuroimaging studies suggests that cortical structures contribute to the control of simple, steady-state human gait (Fukuyama et al. 1997; Gwin et al. 2011; la Fougère et al. 2010; Miyai et al. 2001; Petersen et al. 2012; Seeber et al. 2014). However, the nature of corticospinal interactions that underlie this control remains poorly understood.

In recent years ambulatory electroencephalography (EEG) has been increasingly used to investigate cortical and corticospinal sensorimotor processes during walking in humans (Artoni et al. 2017; Bradford et al. 2016; Bruijn et al. 2015; Bulea et al. 2015; Gwin et al. 2011; Knaepen et al. 2015; Luu et al. 2017; Oliveira et al. 2017b; Petersen et al. 2012; Seeber et al. 2014; Seeber et al. 2015; Severens et al. 2012; Sipp et al. 2013; Storzer et al. 2016; Wagner et al. 2016; Wagner et al. 2012; Wagner et al. 2014; Winslow et al. 2016). Time-frequency analysis revealed that cortical oscillations and corticospinal interactions are modulated relative to the gait cycle at theta (4-7 Hz), alpha (8-12 Hz), beta (13-30 Hz) and gamma (>30 Hz) frequencies. However, there are considerable discrepancies in the precise temporal and spectral patterns of cortical dynamics, probably due to the wide variety of walking tasks used (e.g. treadmill walking at different speeds, on gradients, with additional gait-stability-challenging tasks, with robotic assistance). For instance, beta oscillations were found to be enhanced during the double support phases of the gait cycle (event-related synchronization, ERS) and to be suppressed during the swing and single support phases (event-related desynchronization, ERD) (Artoni et al. 2017; Bradford et al. 2016; Bruijn et al. 2015; Bulea et al. 2015; Cheron et al. 2012; Gwin et al. 2011; Knaepen et al. 2015; Severens et al. 2012). In contrast, other groups observed ERS at 24-40 Hz (low gamma) during early and mid-swing, and ERD towards the end of the swing phase and during double support (Seeber et al. 2014; Seeber et al. 2015; Storzer et al. 2016; Wagner et al. 2012; Wagner et al. 2014).

There are also some initial reports which suggest that these cortical oscillations are transmitted to spinal motoneurons during walking in humans (Artoni et al. 2017; Brantley et al. 2016; Petersen et al. 2012; Winslow et al. 2016). In nine participants, Petersen *et al.* (2012) showed corticomuscular coherence (CMC) between electroencephalographic (EEG) and electromyographic (EMG) signals from the tibialis anterior muscle (TA) at 8–12 Hz and 24–40 Hz during walking. CMC was observed approximately 400 ms prior to heel strike at two different walking speeds (1 km h^−1^ and 3.5–4 km h^−1^). It is uncertain which exact phase of the gait cycle this would correspond to, as 400 ms before heel strike would coincide with different phases of the gait cycle at these two walking speeds (Castermans and Duvinage 2013). The authors suggested that CMC at 24-40 Hz during walking represents an efferent drive from cortex to spinal motoneurons, as they observed a negative imaginary part of coherency indicating a positive time lag from EEG to EMG (Petersen et al. 2012); however, the absolute time lag was not investigated. Recently, Artoni and colleagues (2017) reported a descending connectivity from motor cortex to leg muscles during undemanding, steady-state treadmill walking, which was strongest for muscles of the swing leg but also present for muscles of the stance leg.

The vast majority of previous studies investigating gait-related cortical oscillations have been conducted on standard, motorized treadmills. Overground walking, however, has been shown to differ from treadmill walking dynamically and mechanically (Alton et al. 1998; Arsenault et al. 1986; Carpinella et al. 2010; Chiu et al. 2015; Dingwell et al. 2001; Lee and Hidler 2008; Ochoa et al. 2017; Riley et al. 2007; White et al. 1998). For example, treadmill walking has been found to reduce kinematic variability (Chiu et al. 2015; Dingwell et al. 2001), increase local dynamic stability (Dingwell et al. 2001), affect inter-limb coordination (Carpinella et al. 2010), modify lower limb muscle activation patterns, and joint moments and powers (Lee and Hidler 2008). These observations suggest that sensorimotor control of locomotion may vary between these two gait modalities, potentially affecting processes at the level of the cortex.

In the present study, we investigate cortical sensorimotor oscillations and corticospinal interactions during treadmill and overground walking in a large sample of healthy, young adults. By examining changes in cortical power and CMC during the gait cycle, we investigate the temporal and spectral profile of corticospinal interactions during human walking. In addition, we investigate inter-trial coherence (ITC; Delorme and Makeig 2004) to assess the phase dynamics associated with the changes in spectral power. By comparing power and phase changes we can distinguish whether event-related components result from an evoked activation that is superimposed on the ongoing background activity (hence, an additive response independent from ongoing activity; evoked response), or whether ongoing activity is altered by means of changes in amplitude and/or phase (i.e. corticospinal interactions modulate the phase of ongoing activity; induced response) (Boonstra et al. 2006; Makeig et al. 2004). Finally, the ‘constant phase shift plus constant time lag model’ (Mima and Hallett 1999) was used to estimate the time lag from the CMC phase spectra and investigate the time lag of corticospinal interactions. Together, these spectral analyses will elucidate the nature of corticospinal dynamics involved in human gait.

## Materials and Methods

Twenty-four healthy young adults (mean (SD), age 25.9 (3.2) years, height 170.4 (9.5) cm, weight 68.8 (12.1) kg; 12 men and 12 women) participated in the study. All experimental protocols were approved by the Human Research Ethics Committee of Queensland University of Technology (#1300000579) in accordance with the Declaration of Helsinki, and all participants gave written informed consent prior to participation.

### Experimental protocol

Participants performed 12-14 minutes of overground walking and approximately seven minutes of treadmill walking at their preferred walking speed (3.3 – 4.8 km h^−1^). The order of treadmill and overground walking was randomised across participants. The first two minutes of each trial served as familiarisation period and was not included in the analysis. In the overground condition, participants walked back and forth along a straight path (approximately 14 m) on a firm surface in the gait laboratory and turned at each end of the room. The turning sections, including acceleration at the beginning of the straight-line path and deceleration at the end (approximately 2.4 m), were labelled by the examiner and not included in further analyses, leaving 8.9 m of straight-line walking for further analysis. In order to obtain a similar amount of straight-line walking data in the overground condition, participants were required to walk for a longer duration (12-14 min) than in the treadmill condition (7 min). Treadmill walking was performed on a conventional treadmill (Nautilus, Pro Series). Both walking conditions were completed barefoot. Participants were instructed not to blink excessively, to reduce swallowing and to relax their face, jaw, and shoulder muscles during the data acquisition periods in order to minimise EEG artefacts. Participants rested for five minutes between overground and treadmill walking in order for residual walking-specific after-motion-effects to dissipate and to prevent fatigue.

Gait velocity was individually matched for treadmill and overground walking. To this end, participants completed a ‘pre-trial’ (overground) of approximately three minutes before the actual experiment commenced in order to determine their preferred walking speed. Their mean walking velocity during the pre-trial was used to set the treadmill belt speed and to monitor and maintain the participants’ walking speed consistent during the overground walking condition. If the participant did not comply with the pre-determined mean walking speed during the overground condition, they were instructed to walk faster/slower.

### Data acquisition

Paired bipolar surface EMG (Ag-AgCl electrodes, 1 cm^2^ recording area, 2 cm between poles) was recorded from the belly of the left and right TA. The TA was chosen as recording site based on previous reports of CMC and intra-muscular coherence during walking (Halliday et al. 2003; Petersen et al. 2012). EMG signals were filtered at 1-1000 Hz. Simultaneous EEG recordings were made using water-based AgCl electrodes placed on the scalp at 10 cortical sites according to the international 10-20 standard (P3, P4, C3, Cz, C4, F7, F3, Fz, F4, F8) using a 24-electrode cap (Headcap for water-based electrodes, TMSi, The Netherlands). Combined mastoids (A1, A2) were used as reference. The ground electrode was situated on the wrist using a wristband (TMSi, The Netherlands). EEG signals were filtered at 1-500 Hz. Footswitches were attached onto the participants’ sole at the heel and the big toe of both feet. Synchronized EEG, EMG, and footswitch signals were recorded with a wireless 32-channel amplifier system (TMSi Mobita, The Netherlands) and sampled at 2 kHz. The recording system and wireless transmitter were placed in a belt bag and tied around the participants’ waist.

### Data analysis

All processing of data and spectral analyses were performed in MATLAB (2017a) using custom written routines.

#### Temporal gait parameters

Temporal gait parameters were extracted from the footswitch recordings. Time points of heel strike and toe-off for both feet were extracted and several gait parameters were calculated including stride time, step time, stride- and step time variability, cadence, swing phase, stance phase, single- and double-support time. Time points of heel strike and toe-off were extracted from the digital output signal of the TMSi Mobita system. The channels of the amplifier system used for the footswitches have a default value of approximately −2.04V when ‘off’ and −1.14V when ‘on’; the threshold for a gait cycle event (heel strike or toe-off) was set to −1.64V. Walking velocity was calculated based on the time it took participants to complete the 8.9 m of straight-line walking during overground walking.

#### Data pre-processing and artefact removal

Prior to spectral analysis, electrophysiological data were pre-processed and artefacts were removed according to the following steps/processes. EEG channels were visually inspected and segments with excessive noise (large-amplitude movement artefacts, EMG activity) were rejected. Next, EEG signals were band-pass filtered (2nd order Butterworth, 0.5-70 Hz) and re-referenced to a common average reference (re-referencing was performed in EEGLAB version 13.6.5, Delorme and Makeig 2004). A common average reference in the post-processing steps has been shown to minimize motion artefacts in EEG signals (Snyder et al. 2015). EEG recordings during overground walking were truncated to the straight-line walking segments (turning segments were removed). Subsequently, independent component analysis (ICA) was performed on the concatenated segments using the infomax ICA algorithm implemented in EEGLAB. Independent components containing eye blink, muscle, or movement artefacts were removed from the data (on average 3.7 components were removed per participant); the remaining components were retained and projected back onto the channels. Components were assessed based on their topographical projections, power spectra and time series. For instance, independent components were classified as ocular artefacts when the topographical map showed a far-frontal projection, the median frequency was below 3 Hz and the time course showed positive peaks lasting a few tenths of a second at ≤3 Hz intervals (Delorme and Makeig 2011).

A bipolar EEG montage was used to assess cortical activity from bilateral sensorimotor cortices: C3-F3 for the left sensorimotor cortex, and C4-F4 for the right sensorimotor cortex (cf. Long et al. 2015). A differential recording between the electrode pairs is most sensitive to activity locally generated between the electrode pairs and suppresses more distant activity. Hence, it can be used to minimize artefacts similar to a common reference approach (e.g. Petersen et al. 2012; Snyder et al. 2015).

EMG data were high-pass filtered (4^th^ order Butterworth, 10 Hz cut-off) and full-wave rectified using the Hilbert transform. There is some discussion on the use of EMG rectification for the estimation of CMC (Halliday and Farmer 2010; McClelland et al. 2012; Neto and Christou 2010). For assessing CMC at low force levels, it has been shown that rectification is appropriate (Boonstra and Breakspear 2012; Farina et al. 2013; Ward et al. 2013). We therefore rectified the EMG signals, as walking qualifies as low force movement. Moreover, rectification of EMG signals in the process of calculating CMC has been applied in previous ambulatory CMC studies (Petersen et al. 2012).

After rectification, EMG signals were demodulated to avoid spurious coherence estimates resulting from periodic changes in EMG amplitude (Boonstra et al. 2009). Walking periodically activates the muscles and results in a rapid increase in EMG amplitude during each gait cycle. These fluctuations in EMG amplitude were removed by demodulating the EMG signal using the Hilbert transform to obtain the instantaneous phase *θ* of the rectified EMG signal. The demodulated EMG signal *y* can be defined as the cosine of the instantaneous frequency or phase *y* = cos{*θ*}, where *y* has amplitude 1 and the same instantaneous phase as the original rectified EMG signal (Boonstra et al. 2009).

#### Spectral analysis

Time-frequency analysis was used to assess changes in corticospinal interactions during walking for two bipolar EEG signals (C4-F4, C3-F3) and rectified and demodulated EMG signals of bilateral TA muscles (TAl, TAr) with respect to heel strike of the left and right foot. To this end, EEG and EMG signals were segmented into *L* = 220 segments of length T = 1000 ms (−800 to +200 ms with respect to heel strike of the right or left foot) for each participant and condition. Heel strike served as reference point (t = 0) in the gait cycle to which all segments were aligned. Event-related power spectra and coherence were computed across the 220 segments using a 375-ms Hanning window and a sliding window method with increments of 25 ms. The window length was determined after visual inspection of different window settings with simulated data and experimental data of a representative participant, showing that a window of 375 ms optimised the trade-off between spectral and temporal resolution.

Time-frequency power is hence defined as:

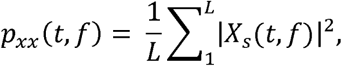

where *X*_*s*_ (*t, f*) is the Fourier transform of the s^th^ segment of signal *x, t* denotes the time relative to the heel strike, *f* the discrete frequency and *L* the number of segments. Likewise, the cross-spectrum is defined as

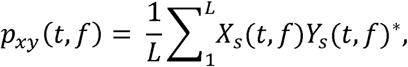

where ‘*’ indicates the complex conjugate. Event-related power was expressed as percentage change from the average by subtracting and dividing by the mean power across the time interval separately for each frequency (Boonstra et al. 2007).

Complex-valued time-frequency coherency is then defined as:

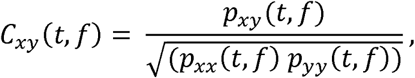

and event-related coherence (Boonstra et al. 2009; Delorme and Makeig 2004) as:

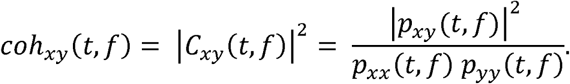

Inter-trial coherence (ITC) can be quantified by averaging the complex-valued Fourier decomposition of *x* over the *L* segments and normalising it by the auto-spectrum:

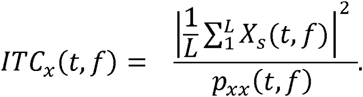

Similar to coherence, ITC is bound between 0 and 1, where 0 indicates the absence of synchronization between the data and the time-locking events and 1 perfect synchronization, i.e., perfect phase reproducibility across trials at a given latency (Delorme and Makeig 2004).

Power and coherence spectra were computed for each participant during overground and treadmill walking, and subsequently averaged across participants to render the grand-average for each estimate and condition. Additionally, event-related EMG envelopes were computed using filtered and rectified EMG data.

To investigate the timing relationship between EEG and EMG activity, we assessed the phase difference between both signals (Halliday et al. 1995). The time lag and phase offset between two signals can be determined from the phase spectra. That is, a constant time lag between two signals results in a linear trend in the phase difference across frequencies (Mima and Hallett 1999). To estimate the time lag from the phase spectra, we fitted a line to the phase spectrum across the frequency range at which significant coherence was observed, and the time lag can be directly calculated from the slope of the regression line (Mehrkanoon et al. 2014; Raethjen et al. 2002). Time lags were separately calculated for different frequency bands and conditions. To this end, complex-valued coherency was averaged across C3F3-TAr and C4F4-TAl to improve the signal-to-noise ratio. Subsequently, the phase spectra were extracted from the averaged coherency estimates, and time lags were calculated for theta, alpha, low beta, high beta and gamma frequency ranges.

## Statistical analysis

Significance level (alpha) was set at 0.05 for all statistical analyses.

Paired sample t-tests (SPSS, version 23) were used to compare gait parameters across conditions (overground, treadmill).

To assess whether the coherence estimates were statistically significant in individual participants, confidence intervals (CI) were constructed based on the number of segments included in the analysis. The 95% confidence interval *CI* was estimated as (Amjad et al. 1997)

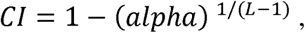

where *alpha* defines the significance threshold and *L* the number of segments. Here *alpha* is set to 0.05 and the resulting CI is hence 1 – (0.05)^1/(220-1)^ = 0.0136. The same CI was also used for ITC.

A two-stage summary statistics approach was used to assess statistical significance of time–frequency coherence at group level. First, magnitude-squared coherence of individual participants was transformed to z-scores using a parametric approach (Halliday et al. 1995). To this end, we first computed the p-values of the coherence values. The p-value of coherence can be estimated as

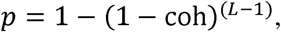

where *coh* is magnitude-squared coherence (Halliday et al. 1995). Then we used the standard normal distribution to convert p-values into z-scores, which have an expected value of 0 and a standard deviation of 1 under the null hypothesis (EEG and EMG signals are independent). Second, a one-sample t-test was used to test whether the individual z-scores were significantly different from zero (Mehrkanoon et al. 2014). This approach was used to test statistical significance of CMC and ITC.

Linear mixed models (LMM) were used to compare the repeated measures of z-transformed coherence across conditions (overground, treadmill). Mean z-transformed coherence over salient time-frequency windows was used for the analysis. Five time-frequency windows were identified: Double support (0 to 125 ms at/after heel strike) at (i) theta (4-7 Hz), (ii) alpha (8-12 Hz), (iii) lower beta (13-20 Hz), (iv) higher beta (21-30 Hz), and (v) gamma frequencies (31-45 Hz). Hence, five separate LMMs were completed (one for each frequency band). Statistically significant main effects were followed up with post-hoc comparisons (t-tests) and family-wise error rate was controlled by using a Bonferroni adjustment for multiple comparisons. The fixed effect factor included in the model was condition; data of both left and right EEG-EMG pairs (C3F3-TAr, C4F4-TAl) were included to compare overground versus treadmill gait. A random effect for participant (i.e. using the participant identification number) was included in the model, which takes into account the repeated coherence measures among participants. The LMM analysis was performed in SPSS (version 23). The same model was used to compare EEG power and ITC across conditions (overground, treadmill). Mean normalized PSD over the salient time-frequency windows per participant was used for the analysis.

An intercept-LMM with a fixed effect factor for condition was used to compare the time lags of the different frequency bands against zero, and across overground and treadmill walking. A significant intercept represents a rejection of the null hypothesis and indicates a non-zero time lag between EEG and EMG signals at group level. A significant effect for condition indicates a difference in time lag between overground and treadmill walking.

## Results

Data from two participants were excluded from the spectral analysis (i.e. for all spectral outcome measures: power, coherence and phase) due to excessive EEG artefacts across all channels and conditions (large-amplitude movement artefacts (>300µV) occurring with regular rhythmicity in each gait cycle throughout the entire record). Another two datasets were excluded during treadmill walking, because fewer than 100 left heel strike triggers were extracted. Thus, group mean estimates for spectral analyses relative to heel strike of the left foot during treadmill walking (i.e. C4F4 for power, and C4F4-TAl for coherence and phase) were based on n = 20 participants. All other spectral estimates (C3F3-TAr coherence/phase and C3F3 power overground and treadmill, C4F4-TAl coherence/phase and C4F4 overground) were based on n = 22 participants.

### Temporal gait parameters

Participants were instructed to walk at their preferred speed, which was matched across both gait conditions. Mean walking speed was not significantly different between overground and treadmill walking (p = 0.36). However, walking speed values were missing for three participants; hence, the group mean estimate was based on n = 19 participants. Table 1 presents all temporal gait parameters during overground and treadmill walking extracted from the footswitch recordings.

**Table 1.** Temporal gait parameters during overground and treadmill walking.

| Gait Parameter | Overground<br>mean (SD)<br>n = 22 | Treadmill<br>mean (SD)<br>n = 22 | T-statistic<br>T(df), p-value |
| --- | --- | --- | --- |
| Walking Speed<br>(km h<sup>-1</sup>) | 4.165 (0.411)* | 4.178 (0.421)* | T(18) = -0.934, p = 0.362 |
| Cadence<br>(strides/min) | 54.088 (3.817) | 55.497 (3.440) | **T(21) = -2.750, p = 0.012** |
| Stride time (s) | 1.115 (0.080) | 1.085 (0.070) | **T(21) = 2.998, p = 0.007** |
| Stride time<br>variability (s) | 0.020 (0.006) | 0.019 (0.019) | T(21) = 0.299, p = 0.768 |
| Step time (s) | 0.551 (0.040) | 0.544 (0.042) | T(21) = 1.273, p = 0.217 |
| Step time<br>variability (s) | 0.011 (0.004) | 0.014 (0.021) | T(21) = -0.752, p = 0.460 |
| Stance phase (s) | 0.696 (0.072) | 0.681 (0.060) | T(21) = 2.043, p = 0.054 |
| Swing phase (s) | 0.407 (0.024) | 0.401 (0.032) | T(21) = 1.337, p = 0.196 |
| Single<br>support (s) | 0.407 (0.024) | 0.401 (0.032) | T(21) = 1.337, p = 0.196 |
| Double<br>support (s) | 0.144 (0.036) | 0.147 (0.046) | T(21) = -0.436, p = 0.667 |

\* n = 19

Most temporal gait parameters were not significantly different between overground and treadmill walking: including duration of step time, step and stride time variability, swing, single and double support phases. However, stride time was significantly longer during overground than during treadmill walking (p = 0.007), and there was a trend towards an extended stance phase during overground walking (p = 0.054). Moreover, participants walked at a higher cadence on the treadmill than they did overground (p = 0.012).

### EMG envelope

Figure 1A and B shows the event-related EMG envelopes for the right and left TA during overground and treadmill walking. The pattern shows increased amplitude during early swing (around −300 ms prior to heel strike), as well as during late swing and early stance (−100 ms prior to +100ms after heel strike). During mid swing (around −200 ms prior to heel strike) the amplitude decreased. The envelope is marked by a double peak at the time point of heel strike (t = 0) when the foot is fully dorsiflexed and at 60 ms after heel strike (early double support). The amplitude increased during early swing already commenced before toe-off of the swing leg. We present single participant data in the appendix.

**Figure 1.**
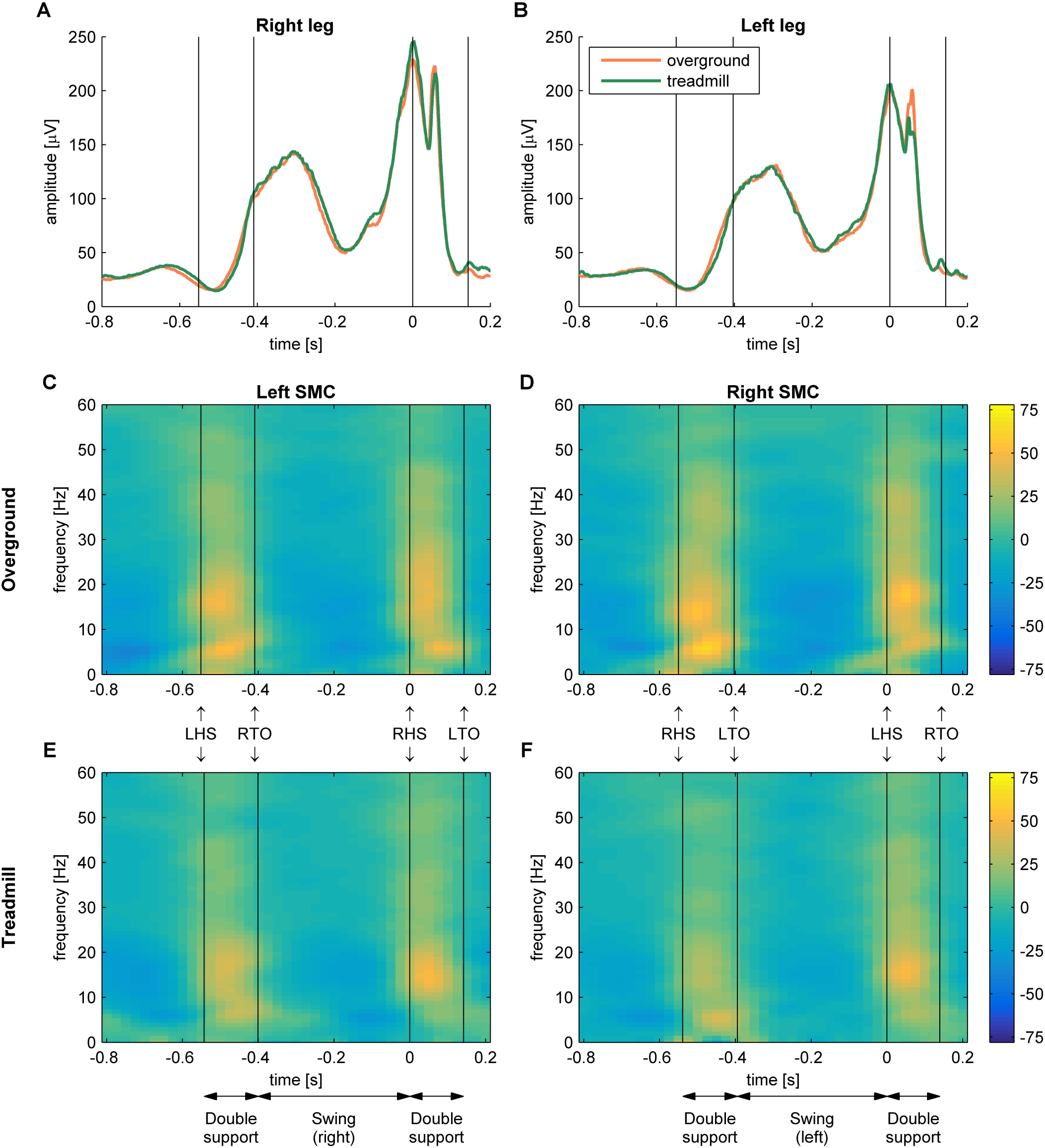
Grand-average EMG envelopes and time-frequency EEG power spectra. EMG envelopes of the right (A) and left (B) TA during overground (orange) and treadmill (green) walking. EEG PSD acquired from bipolar EEG signals of the left sensorimotor cortex (C3-F3) during overground (C) and treadmill walking (E), and of the right sensorimotor cortex (C4-F4) during overground (D) and treadmill walking (F), showing the percent change from the average. X-axis shows time points of gait cycle in seconds, relative to heel strike (t=0) of the left (B, D, F) and right (A, C, E) foot. LHS, left heel strike; LTO, left toe-off; SMC, sensorimotor cortex; RHS, right heel strike; RTO, right toe-off; TA, tibialis anterior.

### Event-related power

The amplitude modulations of cortical activity revealed a different pattern. Increased EEG power was observed across a range of frequencies during double support, while power reduced during the swing and single support phases (Figure 1C to F). The greatest power increase was observed during double support between approximately 15 to 25 Hz, with an increase of up to 55% relative to the average across the whole gait cycle.

The statistical comparison using LMM revealed that EEG power during double support in the theta band was significantly greater during overground compared to treadmill walking (F(1, 63.9) = 5.04, p = 0.03; Figure 2A). The increase in theta power was 29.7% during double support for overground and 15.2% for treadmill walking; the estimated mean difference was 14.5% [95% CI: 1.6-27.4]. No significant differences (p ≥ 0.14) between overground and treadmill walking were observed for the other frequency bands (Figure 2A).

**Figure 2.**
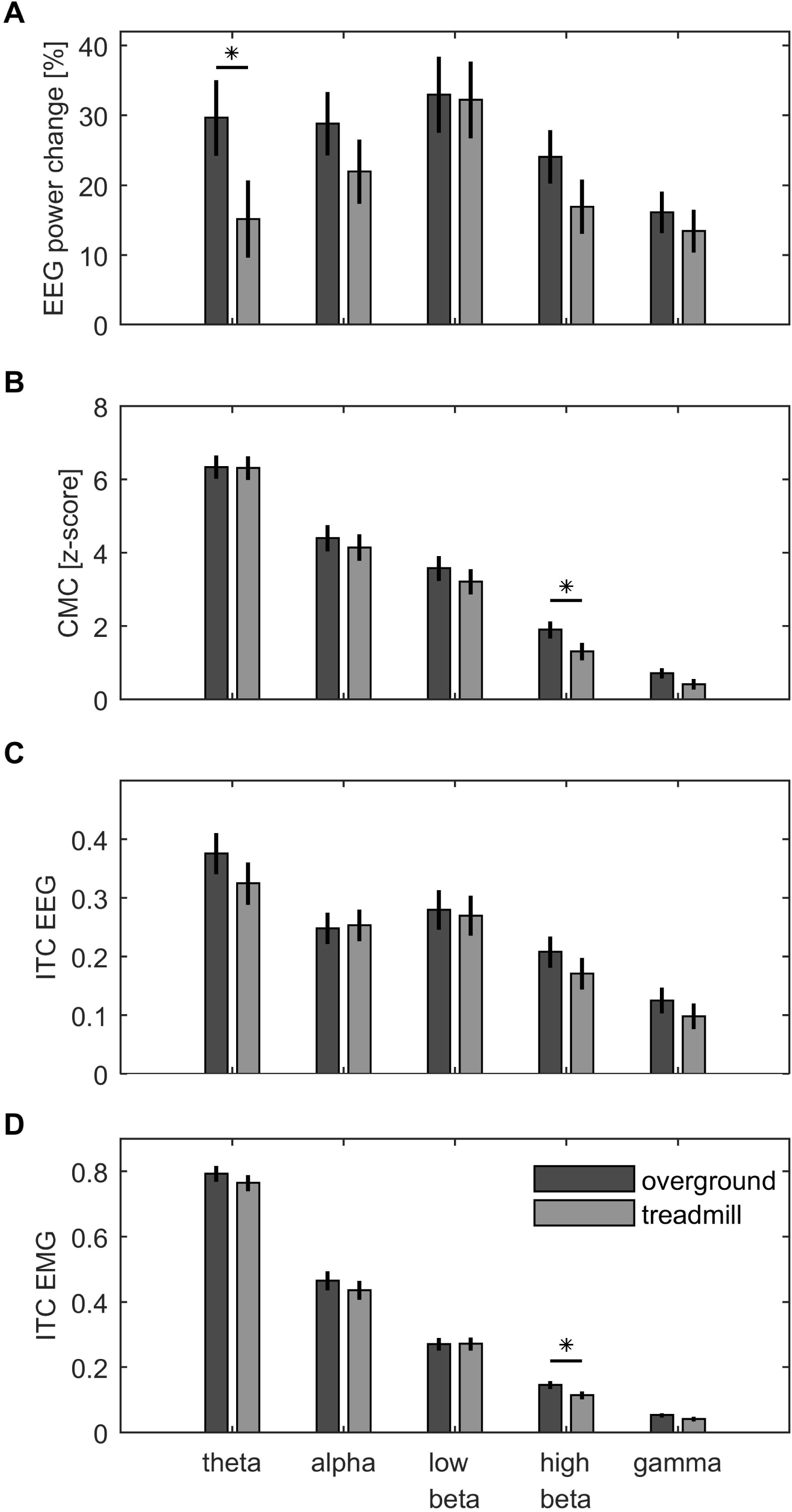
Grand-average EEG power, CMC and ITC during double support. Data during double support (0-125 ms) averaged across sides/channels (C3F3, C4F4 for EEG data; TAl and TAr for EMG data), depicted for each frequency band (theta, alpha, low beta, high beta, gamma) during overground and treadmill walking. Bars show the group mean, vertical lines show two SEM (±1 SEM). Significant differences between overground and treadmill conditions are marked with asterisks. (A) Normalized EEG PSD, (B) z-transformed coherence, (C) EEG inter-trial coherence, (D) EMG inter-trial coherence. CMC, corticomuscular coherence; ITC, inter-trial coherence; TA, tibialis anterior.

### Corticomuscular coherence

During both gait conditions CMC increased during the double support phase of the gait cycle at frequencies between 0 to approximately 45 Hz and little or no CMC was observed during the swing and single support phase (Figure 3). During the double support phase, CMC decreased at higher frequencies showing the highest CMC for frequencies <10 Hz, medium CMC for frequencies 11-20 Hz, lowest CMC for frequencies >25 Hz. CMC was statistically significant during double support in all frequency bands for both sides/channels and gait conditions (p < 0.02).

**Figure 3.**
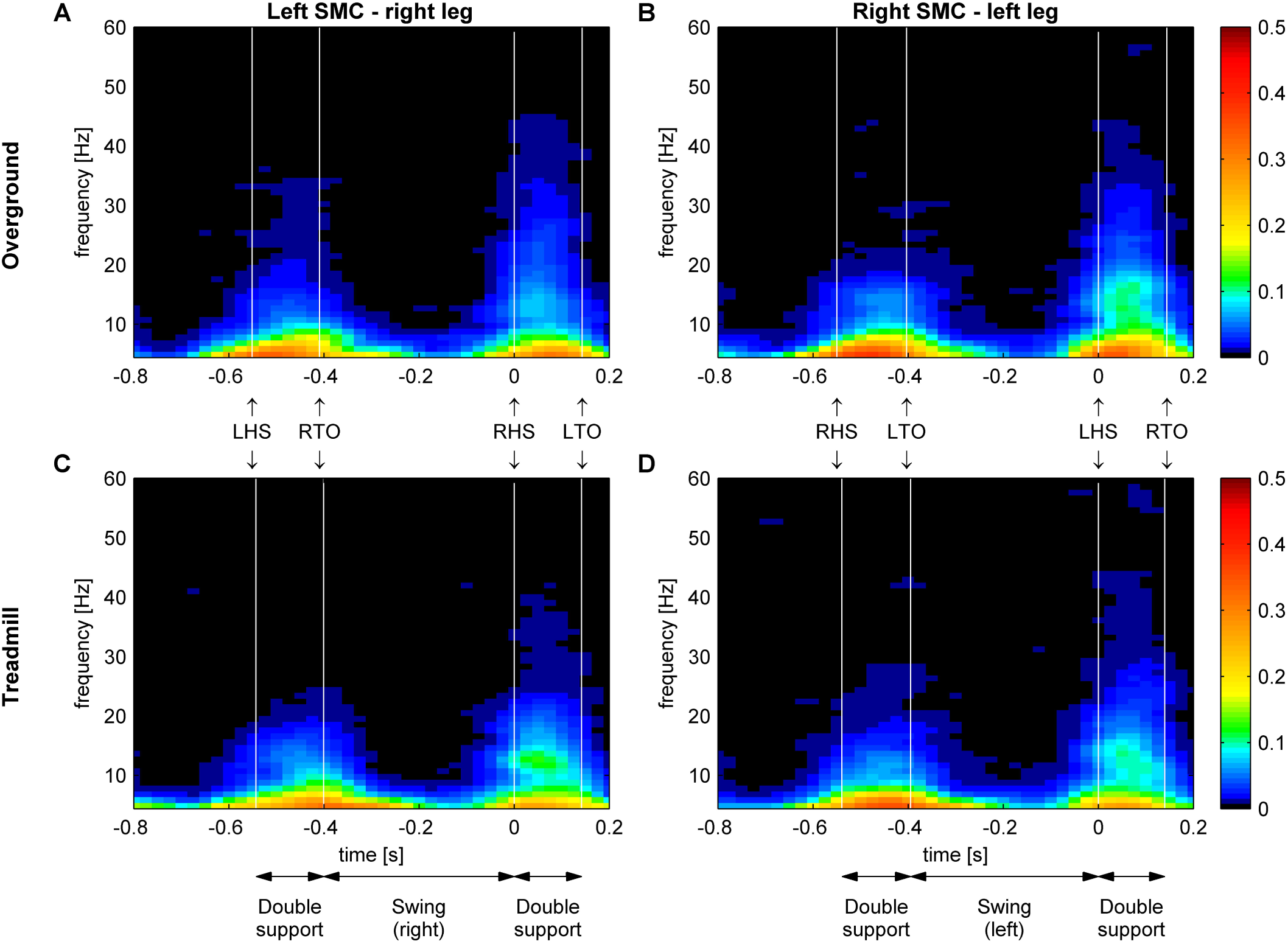
Grand-average time-frequency coherence between EEG and EMG. Corticomuscular coherence between bipolar EEG from the left sensorimotor cortex (C3F3) and EMG from the right TA during overground (A) and treadmill (C) walking; coherence of bipolar EEG from the right sensorimotor cortex (C4F4) and EMG from the left TA during overground (B) and treadmill (D) walking. X-axis shows time points of gait cycle in seconds, relative to heel strike (t=0) of the left (B, D) and right (A, C) foot. Y-axis shows frequencies in Hz. Colour-coding shows the coherence value. LHS, left heel strike; LTO, left toe-off; SMC, sensorimotor cortex; RHS, right heel strike; RTO, right toe-off; TA, tibialis anterior.

The statistical comparison of the gait conditions using LMMs revealed that CMC in the high beta band was significantly greater during overground compared to treadmill walking (F(1, 63.7) = 4.44, p = 0.04; Figure 2B). Grand-average z-transformed high beta coherence was 1.90 during overground and 1.31 during treadmill walking; the estimated mean difference was 0.59 [95% CI: 0.03-1.14]. No significant differences between overground and treadmill walking were observed in the other frequency bands (p ≥ 0.06).

#### Inter-trial coherence

EEG revealed a significant ITC increase during the double support phases of the gait cycle at frequencies between 4 to 50 Hz, and little or no ITC during the swing and single support phases (Figure 4). During the double support phase, ITC decreased at higher frequencies showing the highest ITC for frequencies <10 Hz, medium ITC for frequencies 11-30 Hz, lowest ITC for frequencies >30 Hz. This pattern was similar for overground and treadmill gait, and for left and right sensorimotor cortices. ITC of bilateral EMG revealed a similar pattern (Figure 5): ITC was largest during the double support phases.

**Figure 4.**
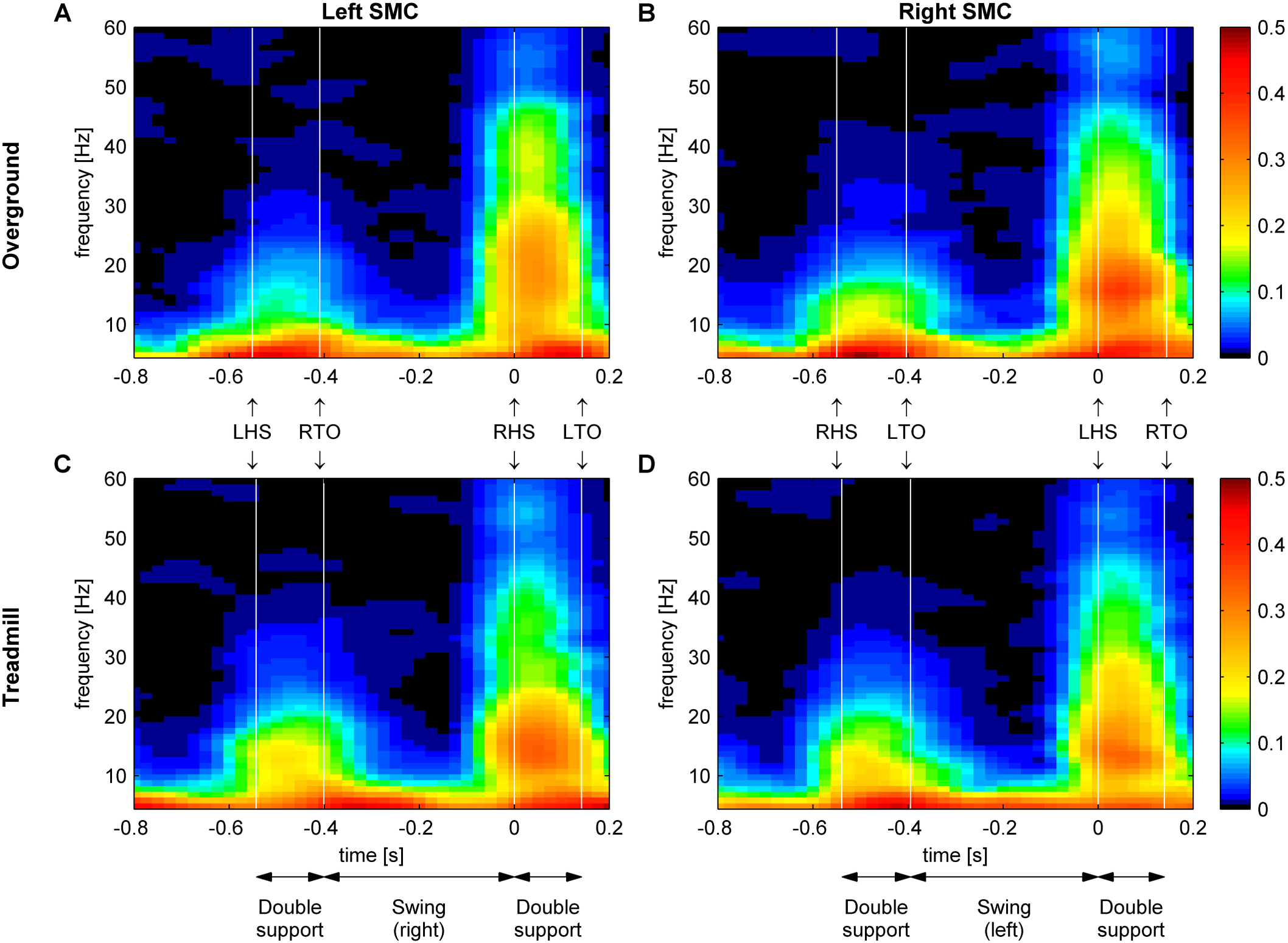
Grand-average time-frequency EEG inter-trial coherence. Inter-trial coherence of the bipolar EEG signals of the left sensorimotor cortex (C3-F3) during overground (A) and treadmill walking (C), and of the right sensorimotor cortex (C4-F4) during overground (B) and treadmill walking (D). X-axis shows time points of gait cycle in seconds, relative to heel strike (t=0) of the left (B, D) and right (A, C) foot. Non-significant values are masked in black. LHS, left heel strike; LTO, left toe-off; SMC, sensorimotor cortex; RHS, right heel strike; RTO, right toe-off.

**Figure 5.**
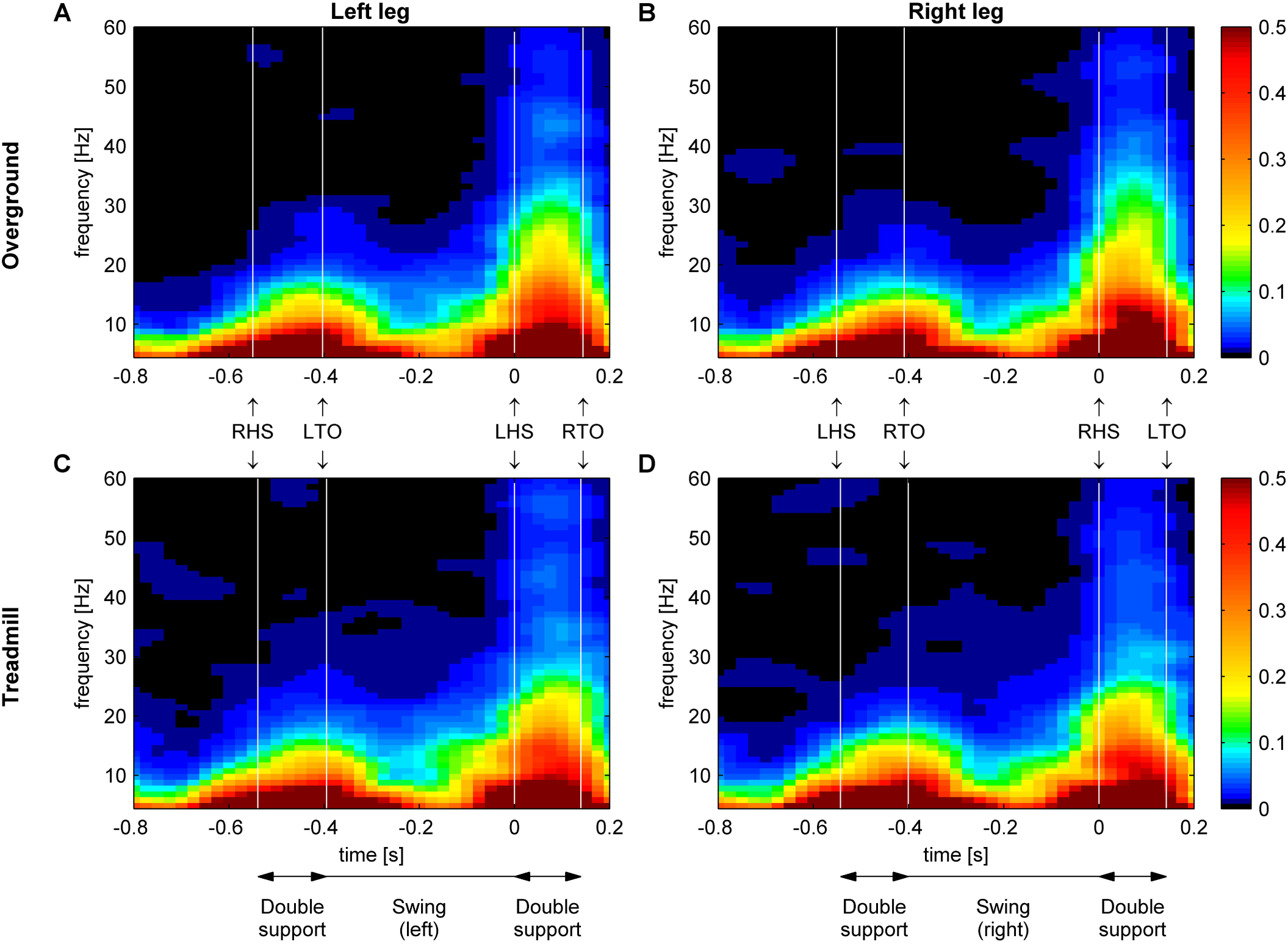
Grand-average time-frequency TA EMG inter-trial coherence. Inter-trial coherence of the EMG signals of the left TA during overground (A) and treadmill walking (C), and of the right TA during overground (B) and treadmill walking (D). X-axis shows time points of gait cycle in seconds, relative to heel strike (t=0) of the left (A, C) and right (B, D) foot. Non-significant values are masked in black. LHS, left heel strike; LTO, left toe-off; RHS, right heel strike; RTO, right toe-off; TA, tibialis anterior.

EEG and EMG ITC was statistically significant during double support in all frequency bands for both sides/channels and gait conditions (p < 0.0001).

The statistical comparison of the gait conditions using LMMs revealed that ITC of the left and right SMC during double support was not significantly different between overground and treadmill walking for any of the frequency bands (p ≥ 0.25; Figure 2C). ITC of the left and right TA was significantly greater during overground compared to treadmill walking in the high beta band (F(1, 63.14) = 7.79, p = 0.007; Figure 2D). Grand-average high-beta ITC was 0.15 during overground and 0.11 during treadmill walking; the estimated mean difference was 0.03 [95% CI: 0.009-0.056]. No significant differences between overground and treadmill walking were observed in the other frequency bands (p ≥ 0.06).

#### Phase spectra

The time delay between EEG and EMG was estimated during overground and treadmill walking separately in each frequency band (theta, alpha, lower beta, higher beta, gamma; Figure 6). We extracted the coherence spectrum in the middle of the double support phase (t = 50 ms) and determined for each participant the frequency bins at which CMC was significant. A linear line was fitted to these frequency bins of the corresponding phase spectra and the time lag was estimated from the slope of the fitted line (see Figure 6A and B for an example in the lower beta band).

**Figure 6.**
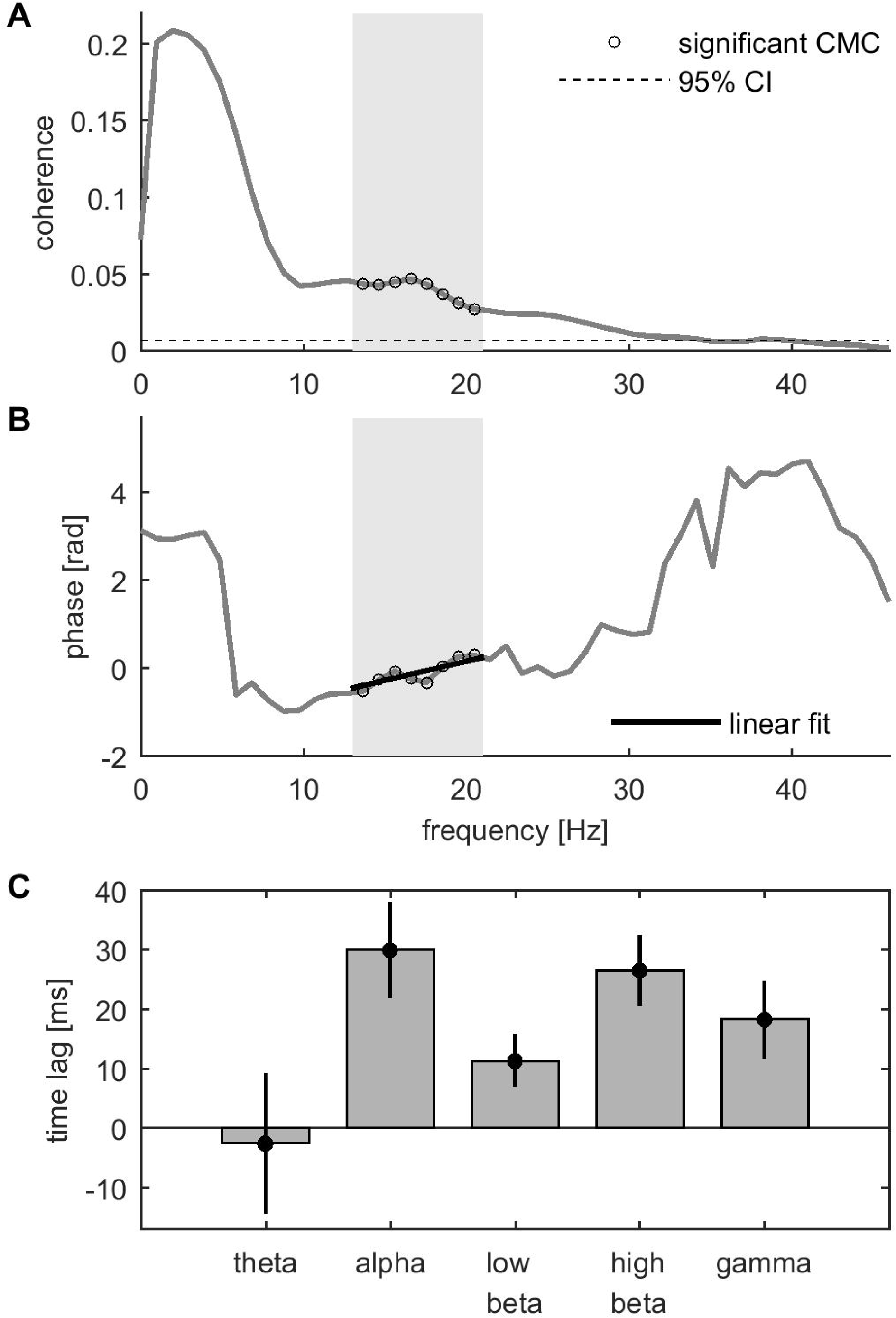
Estimation of time lag between EEG and EMG. (A) Corticomuscular coherence between bipolar EEG from the sensorimotor cortex and contralateral TA EMG at the time point +50 ms (mid double support) during overground walking. Data were averaged across sides/channels (C3F3-TAr, C4F4-TAl). Grey patches indicate low beta frequency range (13-21 Hz), dots depict FFT frequency bins with significant CMC, horizontal dashed line shows upper limit of 95% CI. (B) Corresponding phase spectrum and fitted regression line in the lower beta band. (C) Estimated mean time lag between EEG and EMG in each frequency band (averaged across overground and treadmill, as statistical analyses identified no significant effect of gait condition on time lag). Bars and dots show the group mean; vertical lines depict the standard error of the mean. Time lags estimated based on slope of fitted regression line of individual phase spectra in each frequency band and condition. Positive time lag values indicate a time lag from EEG to EMG. CI, confidence interval; CMC, corticomuscular coherence.

Statistical analyses of time lag revealed that time lag during double support was greater than zero in the alpha (F(1, 33) = 13.3, p = 0.0009), low beta (F(1, 39) = 6.88, p = 0.012), high beta (F(1, 25) = 19.4, p = 0.0002), and gamma bands (F(1, 17) = 7.84, p = 0.012), but not in the theta band (p = 0.84). This indicates that EEG signals were leading EMG signals in alpha, beta and gamma frequency bands. The grand-average time lag was 30.0 ms [95% CI: 13.3-46.8] in the alpha band, 11.4 ms [95% CI: 2.6-20.8] in low beta, 26.5 ms [95% CI: 14.1-38.9] in high beta, 18.3 ms [95% CI: 4.5-32.0] in the gamma band (Figure 6C). No significant differences in time lag were observed between overground and treadmill walking (p ≥ 0.15).

## Discussion

Spectral measures (power, CMC and ITC) were examined during overground and treadmill walking to investigate corticospinal dynamics during both gait conditions. Significant CMC at theta, alpha, beta and gamma frequencies (4-45 Hz) was found between the sensorimotor cortex and the contralateral TA muscle during the double support phase. CMC was largely absent during the swing phase. Similarly, EEG power and ITC increased in these frequency bands during double support and decreased during swing for both gait conditions. CMC and ITC of EMG was significantly enhanced in the high beta band during overground compared to treadmill walking, as well as EEG power in the theta band. Alpha, beta and gamma CMC during double support showed a positive time lag from EEG to EMG, suggesting that cortical activity was leading muscle activity. Most temporal gait parameters were not significantly different between overground and treadmill walking, except stride time (longer overground) and cadence (lower overground).

### Phasic activity during double support

For both overground and treadmill walking, we found that EEG power, CMC and ITC increased during the double support phase of the gait cycle and decreased during swing/single support. Cortical and corticospinal synchronization increased at a broad frequency range, including theta, alpha, beta and gamma frequencies. These results highlight that cortical and corticospinal activity is modulated dependent on the phase of the gait cycle and suggest that the cortex is intermittently involved in controlling human gait.

The cortical power fluctuations during overground and treadmill walking observed in this study closely resemble those previously reported in ambulatory EEG studies conducted on a treadmill. A number of studies found increased power over sensorimotor cortical areas at the end of stance and during double support, and decreased power during the swing/single support phase (Artoni et al. 2017; Bradford et al. 2016; Bruijn et al. 2015; Bulea et al. 2015; Cheron et al. 2012; Gwin et al. 2011; Knaepen et al. 2015; Luu et al. 2017; Oliveira et al. 2017b; Severens et al. 2012). Notably, one previous study (Bulea et al. 2015) reported similar changes in cortical power to those in the current study during walking on a user-driven treadmill, a condition which simulates overground walking more closely than walking on a standard, motorized treadmill. However, other studies reported increased power during the swing phase and reduced power during double support (Seeber et al. 2014; Seeber et al. 2015; Storzer et al. 2016; Wagner et al. 2012; Wagner et al. 2014). Walking tasks in these studies included treadmill walking in a robotic driven gait orthosis (Seeber et al. 2014; Seeber et al. 2015; Wagner et al. 2012; Wagner et al. 2014) and overground walking (Storzer et al. 2016). While robot assisted walking is quite different from steady-state treadmill walking, which may help to explain discrepant findings, the changes in cortical power found during overground walking by Storzer *et al.* (2016) differ substantially from our results. In their study, participants were required to walk at a cadence of 40 strides per minute (although no audio-visual cues were provided to ensure compliance) and walking speed was not kept constant throughout the trial, whereas our participants chose their own cadence for each gait condition (approximately 55 strides per minute) and walking speed was kept constant (∼4 km h^−1^).

The time-frequency profiles of CMC largely match the profiles of cortical oscillations. To our knowledge, only one study (Petersen et al. 2012) and two case-reports (Brantley et al. 2016; Winslow et al. 2016) have previously investigated CMC during walking. Petersen et al. (2012) found significant CMC between motor cortex and TA at 8-12 and 24-40 Hz during treadmill walking, which partly overlap with the frequency ranges in the current study. While all studies show corticospinal interactions during steady-state walking in humans, the temporal profiles of CMC may differ: We found CMC mainly during double support, whereas Petersen et al. (2012) observed CMC ∼400 ms prior to heel strike at two different walking speeds (1 km h^−1^ and 3.5^−4^ km h^−1^). It is uncertain which phase of the gait cycle this exactly corresponds to, as 400 ms before heel strike would coincide with different phases of the gait cycle at these two walking speeds (Castermans and Duvinage 2013). A systematic investigation of CMC at different walking speeds may help to resolve the timing of intra-stride modulations of CMC, as the activations patterns of the TA muscle are strongly modulated by walking speed (den Otter et al. 2004). The case-reports by Winslow et al. (2016) and Brantley et al. (2016) investigated CMC during overground walking in a single subject over a single gait cycle: Winslow et al. (2016) observed CMC between motor cortex and TA at approximately 18-23 Hz just after heel strike and just after toe-off, whereas Brantley et al. (2016) reported CMC between motor cortex and TA at frequencies below 5 Hz throughout the gait cycle. Artoni and colleagues (2017) recently showed a descending connectivity from motor cortical regions to different leg muscles during the swing phase of steady-state treadmill walking. This unidirectional, top-down connectivity was strongest for muscles of the swing leg but also present for muscles of the stance leg during the swing phase. Corticomuscular connectivity was stronger for the TA muscles (of both swing and stance leg) than for other more proximal muscles. However, corticomuscular connectivity was only assessed during the swing phase but not during other phases of the gait cycle (e.g. stance or double support).

The TA EMG envelopes we present in Figure 1A and B (and in Figure 7 in the Appendix) are consistent with previous reports in the literature (Campanini et al. 2007; Courtine and Schieppati 2003; Halliday et al. 2003; Perry 1992; Sutherland 2001; Yang and Winter 1984). Overall, the temporal profiles of our EMG envelopes reveal a well-identified double burst in which the TA is first activated during the onset of swing and then again at the end of swing around heel strike. At the time of heel strike the TA is maximally dorsiflexed within the gait cycle, and after heel strike the TA is performing an eccentric contraction (plantarflexion) which controls foot drop. As the increase of CMC during double support overlaps with the second burst of TA activity, CMC may potentially be involved in maintaining balance during gait by controlling plantarflexion after heel strike. It is interesting that in hemiplegic subjects tibialis anterior activity during swing phase was not affected while they tended to lack the normal second peak of activity at initial foot contact (Burridge et al. 2001). Further experimental investigation of CMC between cortex and leg muscles other than the TA during walking may help to resolve the functional role of CMC during gait. If CMC reflects the descending drive to the muscle, one would expect to find CMC at different phases of the gait cycle for different leg muscles. However, it is worthwhile noting that during dynamic ankle movements beta CMC has been observed during dorsiflexion for two antagonistic muscles (Yoshida et al. 2017).

**Figure 7.**
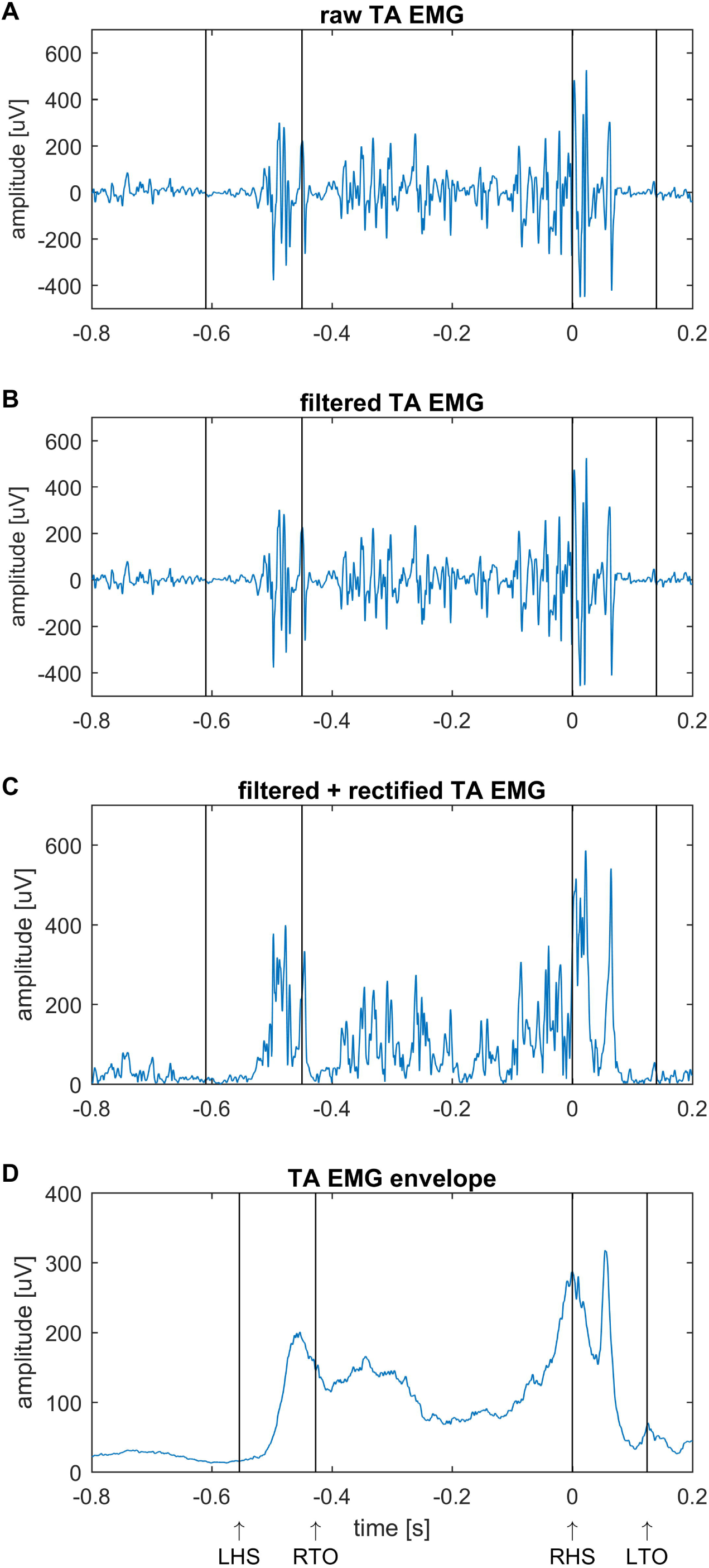
Processing of EMG signals. EMG of right TA of one participant during overground walking. (A) Raw EMG (as recorded) of one step, (B) filtered EMG of one step, (C) filtered and rectified EMG of one step, (D) Mean of 220 steps (envelope) of filtered and rectified EMG. LHS, left heel strike; LTO, left toe-off; RHS, right heel strike; RTO, right toe-off; TA, tibialis anterior.

### Evoked versus induced activity

In addition to the increase in power and CMC, we also observed increased ITC during the double support phases at a broad range of frequencies (theta, alpha, beta and gamma frequencies). ITC quantifies phase locking of the EEG and EMG signals to the time-locking events (Makeig et al. 2004), in this case the heel strike. Phase locking has been widely used to investigate the neural mechanisms that generate event-related brain responses (Fell et al. 2004; Makeig et al. 2004; Shah et al. 2004). Two opposing theories have been discussed that may generate time-locked components: ‘phase resetting’ in which a cortical input resets the phase of ongoing EEG rhythms (induced activity) and ‘evoked responses’ in which input evokes an additive, neural-population response that is independent from ongoing activity. Importantly, induced and evoked activity are not distinguished based on whether the input is generated by a stimulus, but whether it reflects a change in ongoing activity or a response that is linearly added to ongoing activity. A key difference between the phase resetting and evoked models is that an evoked response will be accompanied by an induced increase in power, whereas such an increase is absent in phase resetting (Fell et al. 2004). Although the distinction between evoked and induced activity is not trivial (Mazaheri and Jensen 2010; Ritter and Becker 2009), the parallel increase in power and ITC observed in the current study is most compatible with an evoked response (cf. Tallon-Baudry et al. 1996). Following this interpretation, the time-frequency profile of power and ITC changes would reflect the waveform of the additive response evoked during the double support phase. The waveform of the evoked EMG response can be observed in the EMG envelopes showing a double peak at the time point of heel strike (t = 0) and at 60 ms after heel strike. This corresponds to a 1000/60 = 16.7 Hz oscillation, which is also the frequency at which a peak in ITC is observed (Figure 4).

If the event-related changes indeed reflect an evoked response, this would require a reinterpretation of the observed increase in EEG power during the double support phases. Changes in cortical power are often interpreted as event-related synchronization and desynchronization (ERS/ERD), which is thought to reflect a change in the local synchronization of a neuronal ensemble (Pfurtscheller and Lopes da Silva 1999). As such, ERS and ERD reflect an induced response of ongoing activity rather than an additive response and would hence not be accompanied by phase locked activity (Tallon-Baudry et al. 1996). As the changes in spectral power observed here closely resemble those previously reported in ambulatory EEG studies (Artoni et al. 2017; Bradford et al. 2016; Bruijn et al. 2015; Bulea et al. 2015; Cheron et al. 2012; Gwin et al. 2011; Knaepen et al. 2015; Luu et al. 2017; Oliveira et al. 2017b; Severens et al. 2012), this may require a broader reinterpretation of the cortical dynamics during walking.

Likewise, ITC of EMG and EEG suggests an alternative interpretation of the CMC observed during double support. Conventionally, CMC is thought to reflect corticospinal synchronization, where corticospinal interactions synchronizes the activity in the cortex and spine through efferent and afferent pathways (Conway et al. 1995; Petersen et al. 2012). However, the observed ITC shows that cortical and spinal activity is phase locked to an external event, the heel strike, and as such may not reflect true corticospinal synchronization. Synchronization requires two self-sustained oscillators that adjust their rhythms due to weak interactions (Pikovsky et al. 2003). Instead, the current results suggest that comparable responses are evoked at the cortical and spinal level (i.e. represent time-locked responses at cortical and spinal level that are added to ongoing activity). Importantly, these responses did not occur simultaneously. We observed positive time lags between EEG and EMG activity at alpha, beta and gamma frequencies, which indicated that cortical response occurred before the spinal response.

### Frequency-specific differences between conditions

We found frequency-specific differences in power, CMC and ITC between overground and treadmill walking. Cortical power during double support was significantly increased at theta frequencies during overground compared to treadmill walking. Similarly, CMC and ITC of EMG at high beta frequencies (21-30 Hz) was significantly enhanced during overground walking. In line with previous research, this highlights that neural activity is affected by the walking task that is performed (Bradford et al. 2016; Bruijn et al. 2015; Bulea et al. 2015; Kline et al. 2016; Knaepen et al. 2015; Lisi and Morimoto 2015; Luu et al. 2017; Oliveira et al. 2017b; Sipp et al. 2013; Storzer et al. 2016; Wagner et al. 2016; Wagner et al. 2012; Wagner et al. 2014). For instance, theta ERS in motor cortical regions has been found to be increased during more complex walking tasks, such as walking on a balance beam (Sipp et al. 2013), on a gradient (Bradford et al. 2016), or on an active, user-driven treadmill (Bulea et al. 2015). In the context of the present study, this may suggest that overground walking may be more demanding than walking on a motorized, standard treadmill, and hence be accompanied by increased theta ERS. During overground walking participants have to actively track their walking speed and temporospatial gait patterns themselves rather than simply following the treadmill belt speed, which dictates walking speed and influences temporospatial parameters externally. This would indeed be supported by our finding that treadmill walking decreased stride time and increased cadence. Theta ERS has also been linked to critical time points of the gait cycle, such as transitions between stance and swing, as a control strategy of postural stability (Bradford et al. 2016; Sipp et al. 2013). In that sense, the differences in theta ERS we found between overground and treadmill walking may be related to a difference in timing during overground walking.

High beta CMC was significantly enhanced during overground walking compared to treadmill walking. To our knowledge, no previous study has compared corticospinal dynamics between locomotor tasks, but only investigated it during treadmill walking (Artoni et al. 2017; Petersen et al. 2012; Winslow et al. 2016). However, a number of studies have assessed cortical oscillations during various gait tasks and have found task-dependent differences in cortical beta power (Bruijn et al. 2015; Bulea et al. 2015; Knaepen et al. 2015; Lisi and Morimoto 2015; Oliveira et al. 2017b; Sipp et al. 2013; Wagner et al. 2016; Wagner et al. 2012; Wagner et al. 2014). These studies have suggested that cortical beta oscillations may be related to controlling gait stability (Bruijn et al. 2015; Sipp et al. 2013), gait adaptations (i.e. step lengthening and shortening, Wagner et al. 2016), sensory processing of the lower limbs (Wagner et al. 2012), visuomotor integration (Oliveira et al. 2017b; Wagner et al. 2014), and speed control (Bulea et al. 2015; Lisi and Morimoto 2015). In our experiment, participants were required to perform steady-state overground and treadmill walking at the same speed and without any perturbations that would have triggered deliberate stepping adaptations. Hence, different levels of high beta CMC during overground and treadmill walking may reflect differences in gait stability, step adaptations, sensorimotor processing and/or speed control in each gait condition in our study.

If the spectral changes reflect an evoked response as suggested above, the time-frequency profile of power and ITC changes may reflect the waveform of the additive response rather than a change in cortical oscillations. High beta ITC of the TA muscle was significantly enhanced during overground compared to treadmill walking, while no differences in cortical ITC was observed. Hence, the observed differences between treadmill and overground walking may reflect a change in the evoked response in the TA muscle. Possibly, changes in the evoked TA response may be related to peripheral mechanisms involved in stride time or cadence, as these parameters were significantly different between treadmill and overground walking in the present study. This interpretation is supported by studies that showed biomechanical differences between overground and treadmill gait (Carpinella et al. 2010; Chiu et al. 2015; Dingwell et al. 2001; Lee and Hidler 2008; Ochoa et al. 2017). In line with this, these potential changes in evoked TA responses could be due to differences in background electrical activity of dorsal horn spinal neurons. In cats, for example, it has been shown that amplitude fluctuations of somatosensory evoked field potentials are positively correlated with spontaneous activity of dorsal horn spinal cord neurons (Manjarrez et al. 2002a; Manjarrez et al. 2002b).

### Functional implications

We found positive time lags between coherent EEG and EMG activity at alpha, beta and gamma frequencies, which was consistent for overground and treadmill walking. A positive time lag is consistent with efferent descending activity and hence provides further evidence for efferent cortical control during walking in humans (Artoni et al. 2017; Petersen et al. 2012). The estimated time lags varied from approximately 12 ms in the lower beta band to 30 ms in the alpha band. The time lag at low beta frequencies (∼12 ms) is consistent with previous studies investigating beta-band CMC in upper limb muscles during isometric contractions, e.g. 11 ms (Mehrkanoon et al. 2014), 15.9 ms (Mima et al. 2000), 9.3 ms (Gerloff et al. 2006) and 7.9 ms (Witham et al. 2011). However, this time lag is shorter than the corticospinal conduction time from the cortex to the TA measured by stimulation (roughly 27 ms, Gross et al. 2000; Petersen et al. 2001; Rothwell et al. 1991; Schubert et al. 1997). Indeed, the time delay estimated from phase spectra appears to underestimate the actual physiological corticospinal conduction times (Witham et al. 2011). Hence, while the absolute time lag found in the current study may be an underestimation of the actual conduction time, a positive time lag was robust at the group level and consistent with a cortical drive to the TA muscle during overground and treadmill walking. Although the time lag from cortex to muscle may be consistent with efferent activity, the temporal precedence does not demonstrate that CMC was generated by the influence that the cortex exerts over the spinal motor pool. That is, interpretations in terms of effective connectivity require an explicit model of that influence (Friston 2011).

By estimating the time lag of coherent activity, the approach assesses the temporal precedence between signals similar to other methods of directed functional connectivity (Friston et al. 2013). It rests on the assumption that dependencies reflect an underlying dynamical process, in which causes precede consequences. That is, neuronal interactions are based on synaptic connections and hence involve propagation delays. However, temporal precedence does not necessarily imply a causal influence between the two evaluated channels (Kaminski et al. 2001; Seth 2010). It is possible that the effect is mediated by another channel or by another group of channels, or by variables that are not included in the measurements. This caveat is particularly relevant in the current study, in which the results of time-frequency results are consistent with an evoked response. It is likely that the cortical and spinal input underlying the evoked responses are partly generated by other neural systems that were not measured in the current study. The interpretation of the observed time lag in terms of efferent and afferent processes should hence be made with caution.

### Limitations

Ambulatory EEG is susceptible to movement artefacts and caution should hence be exerted when interpreting phasic changes in EEG activity (Castermans et al. 2014; Costa et al. 2016; Gwin et al. 2010; Kline et al. 2015; Nathan and Contreras-Vidal 2016; Oliveira et al. 2016; Snyder et al. 2015). In this study, we applied multiple strategies to mitigate the effects of motion artefacts: We excluded gait cycles with excessive artefacts, we performed ICA, and we calculated a bipolar derivative EEG signal, which removes further shared artefactual activity across channels similar to a re-referencing approach (Snyder et al. 2015). This way, a bipolar EEG signal reflects nearby cortical activity that is locally generated between the electrode pairs. With respect to the EMG recordings, we high-pass filtered, rectified, and demodulated the signals in order to remove low-frequency artefacts and periodic amplitude modulations that could distort the coherence estimates.

The parallel increase in power, ITC and CMC is consistent with an evoked response and it is hence possible that these responses are generated by mechanical artefacts. However, the positive time lags in the alpha, beta and gamma bands suggest that the results of this study were not due to conduction of non-physiological artefacts, which would have likely resulted in a zero time lag (Petersen et al. 2012), but reflect genuine neural activity. The time lag between coherent theta activity was not significantly different from zero, which may suggest residual artefact contamination at theta frequencies specifically. Previous studies indeed found that oscillatory activity at low frequencies may be more affected by movement artefacts than at other/higher frequency ranges (Castermans et al. 2014). The event-related power profiles observed in this study are similar to those in multiple previous studies on cortical oscillatory activity during walking (Artoni et al. 2017; Bradford et al. 2016; Bruijn et al. 2015; Bulea et al. 2015; Cheron et al. 2012; Gwin et al. 2011; Knaepen et al. 2015; Severens et al. 2012). This would imply that the observed time-frequency modulations of cortical power and CMC in the current and previous studies reflect genuine neural interactions, or alternatively, are all affected in similar way by non-physiological artefacts. EEG acquisition during dynamic movement remains challenging and methods to minimize movement artefacts in ambulatory EEG is an ongoing pursuit (Kline et al. 2015; Oliveira et al. 2017a; Snyder et al. 2015).

## Conclusion

This study shows coherent corticospinal activity during the double support phase of the gait cycle during steady-state overground and treadmill walking in healthy, young adults. Parallel increases in power and ITC suggest evoked responses at spinal and cortical populations rather than a modulation of ongoing corticospinal interactions. A positive time lag between EEG and EMG signals at alpha, beta and gamma frequencies is consistent with efferent activity, but needs to be interpreted with caution. Moreover, this study shows that high beta CMC and ITC of EMG differed between overground and treadmill walking. This may reflect a change in the evoked response in the TA muscle dependent on the gait modality, possibly indicating different peripheral mechanisms that control stride timing. These findings may help to identify neurophysiological mechanisms that are important for understanding cortical control of human gait in health and disease.

## Appendix

Figure 7 shows EMG data of the right TA of one participant during overground walking. These single participant data are similar to the group average data presented in Figure 1A and B.

## Acknowledgements

We thank Bridie O’Connell and Kathryn McIntosh for support with data collection; Kevin McGill, Steve Mehrkanoon and Christopher Thompson for valuable discussions on the z-transformation of coherence; the QUT HPC and Research Support Group for access to computational resources.

